# Genome-wide diversity of chromosomal inversions and their disease relationships

**DOI:** 10.1101/2025.09.25.678440

**Authors:** Scott C. Sauers, Bailey Huepenbecker, PingHsun Hsieh

**Affiliations:** Bioinformatics and Computational Biology Graduate Program, University of Minnesota, Twin Cities, MN, USA; Department of Genetics, Cell Biology, and Development, University of Minnesota, Twin Cities, MN, USA; Institute for Health Informatics, University of Minnesota, Twin Cities, MN, USA

**Keywords:** Inversion, Recurrent mutation, Evolution, PheWAS, Human disease

## Abstract

Chromosomal inversions shape evolution and are implicated in human disease, yet their effects on genomic variation and health outcomes remain poorly understood. We analyze genome-wide human inversion polymorphisms, contrasting single-event and recurrent loci. Inversion recurrence is validated using structured-coalescent simulations. We show that single-event inversions evolve in near-complete isolation: inverted haplotypes show ∼16-fold lower diversity and strong differentiation from direct haplotypes (median *F*_*ST*_ = 0.33). By contrast, recurrent inversions maintain gene flow, resulting in similar diversity across orientations and ∼4-fold lower differentiation. We further find marked differences in coding sequence conservation between single-event and recurrent inversions. Using the NIH All of Us biobank, we impute inversions and identify four inversions with significant disease associations. Notably, the 17q21 inversion is associated with reduced risk of cognitive decline (OR=0.919) and breast cancer (OR=0.910) but with increased obesity risk (OR=1.097), consistent with pleiotropic selection. These findings establish inversions as major drivers of human genetic diversity and disease, with evolutionary outcomes critically dependent on recurrence.

## Introduction

Inversions are structural mutations that reverse a chromosomal segment along with its gene content and are ubiquitously found across species^1-8^. Recombination between heterokaryotypes (inverted vs. direct orientations) is largely suppressed because chiasmata rarely form correctly, making inversions effective barriers to gene flow. This reduction in recombination increases drift, hitchhiking, and the accumulation of linked alleles^1,3,4^. In primates, thousands of inversions have been identified across great apes, and hundreds remain polymorphic in humans^5,6^. Many human inversions are also linked to diseases including blood disorders, cancer, and neurodevelopmental conditions^5,9,10^, highlighting their importance for health.

The drastically reduced recombination rate between haplotypes of opposite orientations influences many evolutionary processes that shape diversity, including drift and natural selection^11^. A long history of theoretical work has established expectations for the distinctive population genetic signatures of variants within inversions^12,13^. Because of limited gene flow, alleles residing on inverted haplotypes evolve in near isolation from the direct orientation. This in turn leads to reduced diversity within the inverted haplotypes due to a reduction in effective population size^3,14,15^. This suppressed recombination promotes genetic isolation and increases divergence between orientations through the accumulation of mutations, as well as formation of excess linkage disequilibrium among alleles in the locus^13,16^. However, rare double crossovers in heterokaryotypes^17^ and gene conversion events^18^ can still lead to genetic exchange between orientations (i.e., gene flux^15^, with exchange more likely in the central part of the inversion than near the ends ^19^. As a result, divergence is often greatest and diversity lowest at inversion breakpoints^3,15^, a pattern observed in Drosophila melanogaster^20^ and in some human inversions^14^. Inversion breakpoints themselves may be directly subject to selection if they disrupt genes or regulatory elements^21^.

Chromosomal inversions shape phenotype, and as a consequence, evolution, by altering gene order, regulation, and linkage^16^. Inversions may also increase fitness by capturing multiple advantageous alleles and shielding them from recombination^13^. By suppressing recombination, inversions preserve sets of alleles such that local adaptation and even speciation can occur, as favorable allele combinations are protected from genetic shuffling^22-24^. Notable examples include thermal adaptation mediated by an ancient 8 Mb inversion (In(3R)Payne) in *D. melanogaster* ^25,26^ and maintenance of beneficial epistasis in *D. pseudoobscura*^18^. At the same time, recombination suppression reduces the effective population size of inverted haplotypes, allowing weakly deleterious alleles to accumulate^27^ Yet purifying selection can still act within inverted sequences, such as in *Calidris pugnax*, where an ancient inversion supergene persists despite the lethality of being an inversion homozygote, with dN/dS consistent with continuous purifying selection across most genes in the inverted haplotype sequences^27^. Due to the lack of recombination, inversions can act as allelic reservoirs, able to preserve ancient haplotypes deeply divergent from the rest of the population, as observed in *Drosophila*^12^.

Detecting inversions in primates, especially humans, has been proven challenging because over half are embedded within large, nearly identical segmental duplications^5,7^. Combining short-read Strand-seq with long-read sequencing, Porubsky et al. (2022)^5^ generated the most comprehensive map of human inversion polymorphisms to date, resolving breakpoints and inversion sequences at base-pair resolution in a cohort of diverse populations. They found that 72% of balanced inversions (i.e., inversions without evidence for any duplication or deletion) occur near segmental duplications or retrotransposons, predisposing these loci to non-allelic homologous recombination and making them hotspots for recurrent structural rearrangements, including inversions. Using an evolutionary approach, evidence of recurrence was observed in 32 of 93 inversions, many of which overlap with morbid copy-number variation loci and likely promote duplication diversity, increase mutability, and predispose haplotypes to disease-causing copy-number variations^5^. Despite these insights, to date the population genetic properties of human inversions, including sequence diversity, divergence, recurrence, and phenotypic effects, remain largely unexplored.

Here, we set out to systematically characterize human chromosomal inversions in terms of genetic diversity, differentiation, selection, and phenotypic impact. We first use a large number of structured coalescent simulations to evaluate the statistical power of the evolutionary framework for detecting recurrent inversions proposed in Porubsky et al. (2022)^5^. We then analyze 93 high-quality single-event and recurrent inversion polymorphisms^5^ through statistical modeling and evolutionary analyses to provide the first glimpse of the variation patterns across the inversion landscape. Finally, we integrate these findings with phenotypic data from the NIH *All of Us* biobank to search for phenotypic associations of inversion loci, revealing the role of inversions in human biology and health.

## Results

### Assessment of recurrent inversion detection using coalescent simulations

To better understand the genomic variation patterns and functional implications for single-event as well as recurrent inversions, we first systematically evaluated the previously published evolutionary approach for detecting recurrent inversions ^5^ using simulations (**Methods**). Briefly, we modeled recurrent inversions as structured coalescent processes with limited or no gene flow (**Fig. 1A and 1B**). When an inversion occurs, recombination between different haplotypes is suppressed, creating a genetic barrier that isolates the inverted haplotypes from the ancestral (direct-oriented haplotype) population. This divergence is modeled as a subpopulation with no gene flow, while subsequent inversions are treated as further divergence events. Gene flow resumes between haplotypes with the same orientation, allowing recombination and genetic variation exchange. We evaluated the statistical power of detecting recurrent inversion loci by simulating different model scenarios for single-event and three recurrent inversion events. A wide range of parameter values was considered, including different time depths of the inversion events, recombination rates, and inversion frequencies (**Figs. 1 and S1-S4, Tables S1-S4, Methods**).

**Fig. 1.**
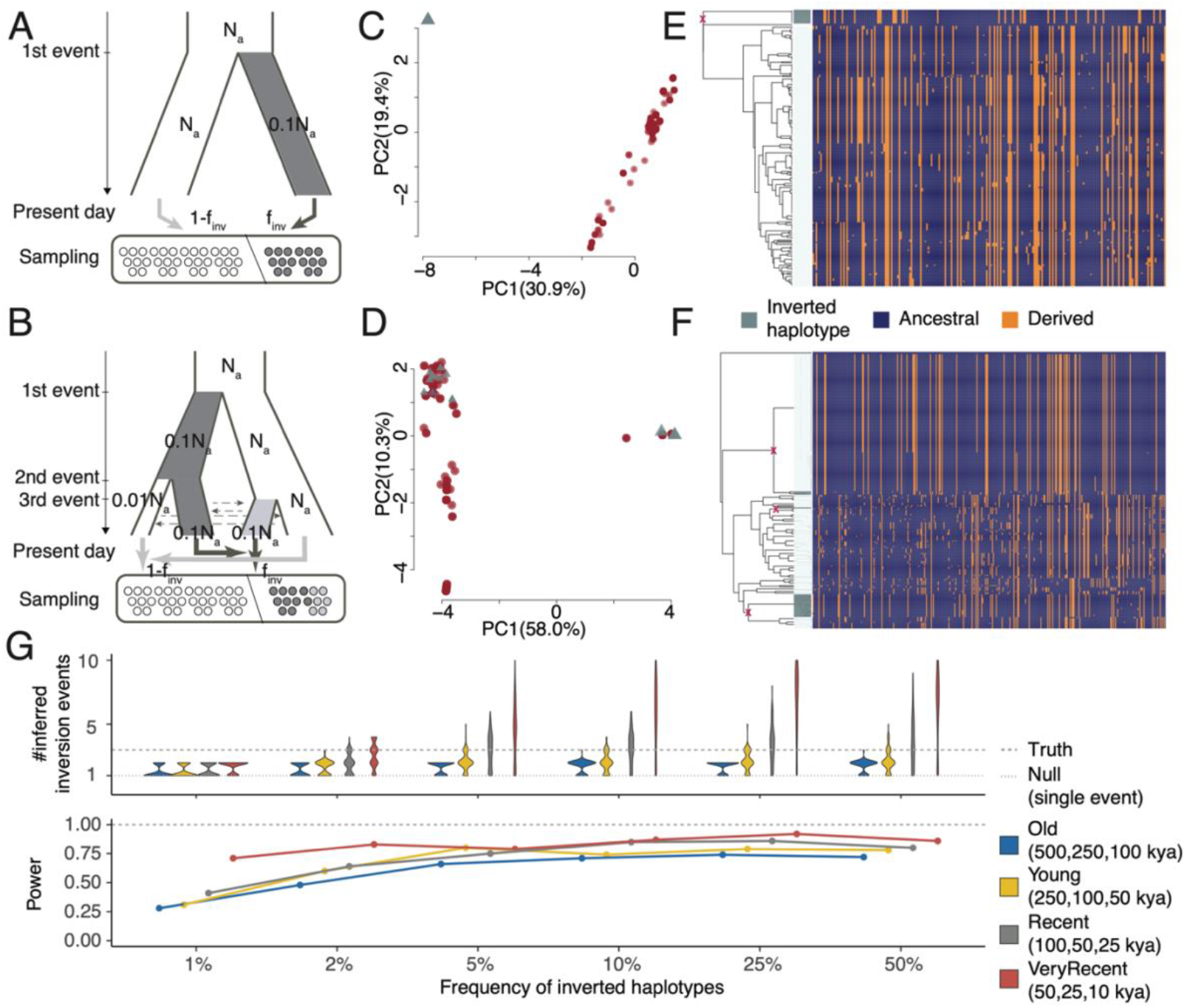
Evaluation and power analysis of the evolutionary approach for detecting recurrent inversions. Schematics of structured coalescent simulations for single-event (**A**) and recurrent (n=3) inversion (**B**). An inversion event is modeled as divergence between inverted (light/dark greys) and direct (white) haplotype populations, followed by isolation without gene flow, reflecting recombination suppression between the populations. N_a_ refers to ancestral population size. Dash arrows indicate gene flows between subpopulations with the same orientation. Each divergence introduces a bottleneck (90% reduction) in the new subpopulation of different orientation from its parental population). Finally, inverted and direct haplotypes are sampled at pre-defined frequencies f_inv_ and 1-f_inv_, respectively, from a random mixture of populations with the same orientation (solid arrows). (**C-F**) Distinct genomic variation patterns between single and recurrent inversion loci. Haplotype-based principal component (PC) analysis indicates unique clusters for inverted (grey triangles) and direct (red circles) in single-event loci (**C**) but not in recurrent (**D**) loci. Percentages on axes indicate the amounts of variance explained by individual PCs. (**E & F**) Dendrograms (centroid hierarchical clustering method) show relationships among inverted and direct haplotypes. Red crosses indicate the inferred inversion events. (**G**) Summary of power analysis for models with three recurrent inversion events across four different time depths and varying inversion frequencies. Each model has 100 replicates. Violin plots (top) show the distribution of inferred inversion events, while curves (bottom) indicate the statistical power to reject the null (single-event) model for each scenario.

Using simulation data, our haplotype-based principal component (PC) and hierarchical clustering-based tree analyses show a clear separation between inverted and direct haplotypes under the single-event models, as opposed to mixing of both haplotypes within the same cluster under the recurrent inversion models (**Figs. 1A-1F**). Across all single-event models considered, our results show that the evolutionary approach achieves low false positive rates (<5%), with the highest rate (4%, 95% C.I.: 0–8%) observed in models of very recent inversion events (50 thousand years ago [kya]) and when recombination rate is high (1 x 10^−6^ per base per generation) regardless of the sampling frequency of inversions. Under the recurrent inversion models, our power to detect recurrent events varies with different scenarios (**Figs. 1G and S1-S4**). In general, the power of identifying recurrent inversion loci decreases when the ages of such events and/or the recombination rate of the loci increase (**Figs. S1-S4**). The strongest factor that affects the power is the sampling frequency of inversions (**Fig. 1G**). The higher the sampling frequency is, the higher the power of the approach is to detect recurrent loci. With moderate recombination rate (1 x 10^−8^ per base per generation), the power ranges from 66–92% when the frequency of inversions is above 5%; however, it decreases to as low as 28% when the inversion frequency is below 5% (**Fig. 1G**). Of note, while the approach in general shows strong power to detect recurrent loci, we found that it tends to overestimate the number of inversion events (**Figs. 1G, S1-S4**), especially for young events. Nevertheless, our power analysis results provide important benchmarks for the evolutionary approach for detecting recurrent inversion loci and strong support for the recurrence classifications of the inversions reported by Porubsky et al. (2022)^5^.

### Differential genomic variation patterns by recurrence and orientation

We set out to characterize the genomic variation of inversion sequences in the human genome using the high-quality sequencing data as well as single-event and recurrent inversion calls derived from a published high-quality call set^5^. This dataset includes 41 unrelated human individuals, from the 1000 Genomes Project panel (82 phased haplotypes). Inversion discovery, phasing, and characterization were performed using a combination of single-cell template strand sequencing (Strand-seq), haplotype-resolved *de novo* genome assemblies generated from long-read sequencing (PacBio HiFi and CLR), and single-molecule optical mapping^5^. The analysis of the present work focused on 93 balanced (**Table S1**) inversions that were confidently called as single-event or recurrent^5^; of which, 32 recurrent balanced inversions, alongside 61 single-event inversions, were identified on autosomal or X chromosomes. Inversions present on the Y chromosome were excluded from all downstream analyses due to the difficulty of variant calling within the sequence. We computed nucleotide diversity (π) and genetic differentiation (*F*_*ST*_) within individual inversion sequences, in contrast to previous studies^5^ focusing on inversions as alleles. We considered these population genetic measurements in relation to (i) orientations (direct vs. inverted haplotypes) and (ii) recurrence of inversions (i.e., single-event vs. recurrent inversions).

We computed π separately in the inverted haplotype group and the direct haplotype group, for each inversion locus with at least two haplotypes present (N=91). Overall, without considering recurrence status, inverted haplotypes had substantially lower diversity (mean π: 0.0002) than direct haplotypes (mean π: 0.0005) (two-sided paired t-test, p=2.1×10^-9^). Within the single-event inversion loci, on average inverted haplotypes had lower diversity (median π: 0.0) than direct-orientation haplotypes (median π: 0.0005); however, we observed an opposite trend within recurrent inversion loci (median π inverted: 0.00046, median π direct: 0.0004). To further investigate these observations, we used a paired linear model which compared differences in π between orientations within each locus to examine the relation of nucleotide diversity to inversion orientation and recurrence status. The model included a recurrence term (single-event vs. recurrent status), and tested the interaction between recurrence and orientation (**Methods**). Within single-event inversions, we observed a significant effect of orientation, with inverted haplotypes showing lower nucleotide diversity compared to direct orientation haplotypes (p = 8.7×10^-21^, two-sided Wald test, **Fig. 2A**). There was no significant difference between orientations within recurrent inversions (two-sided Wald p = 0.489). The model estimated that compared to direct haplotypes, inverted haplotypes had ∼16.4-fold lower nucleotide diversity in single-event inversions, yet without significant differences in diversity among recurrent inversions. Notably, we observed a significant interaction between orientation and recurrence (p = 1.2×10^-19^, Wald test), indicating that the effect of orientation depends on recurrence. That is, the association between inverted orientation and reduced diversity is stronger in single-event inversions (**Fig. 2A**). Indeed, within inverted haplotypes, median nucleotide diversity was markedly higher for recurrent inverted haplotypes (π = 3.98×10^-4^, n = 32) compared to single-event inverted haplotypes, in which the median inversion sequence group contained no variation (π = 0, n = 61).

**Fig. 2.**
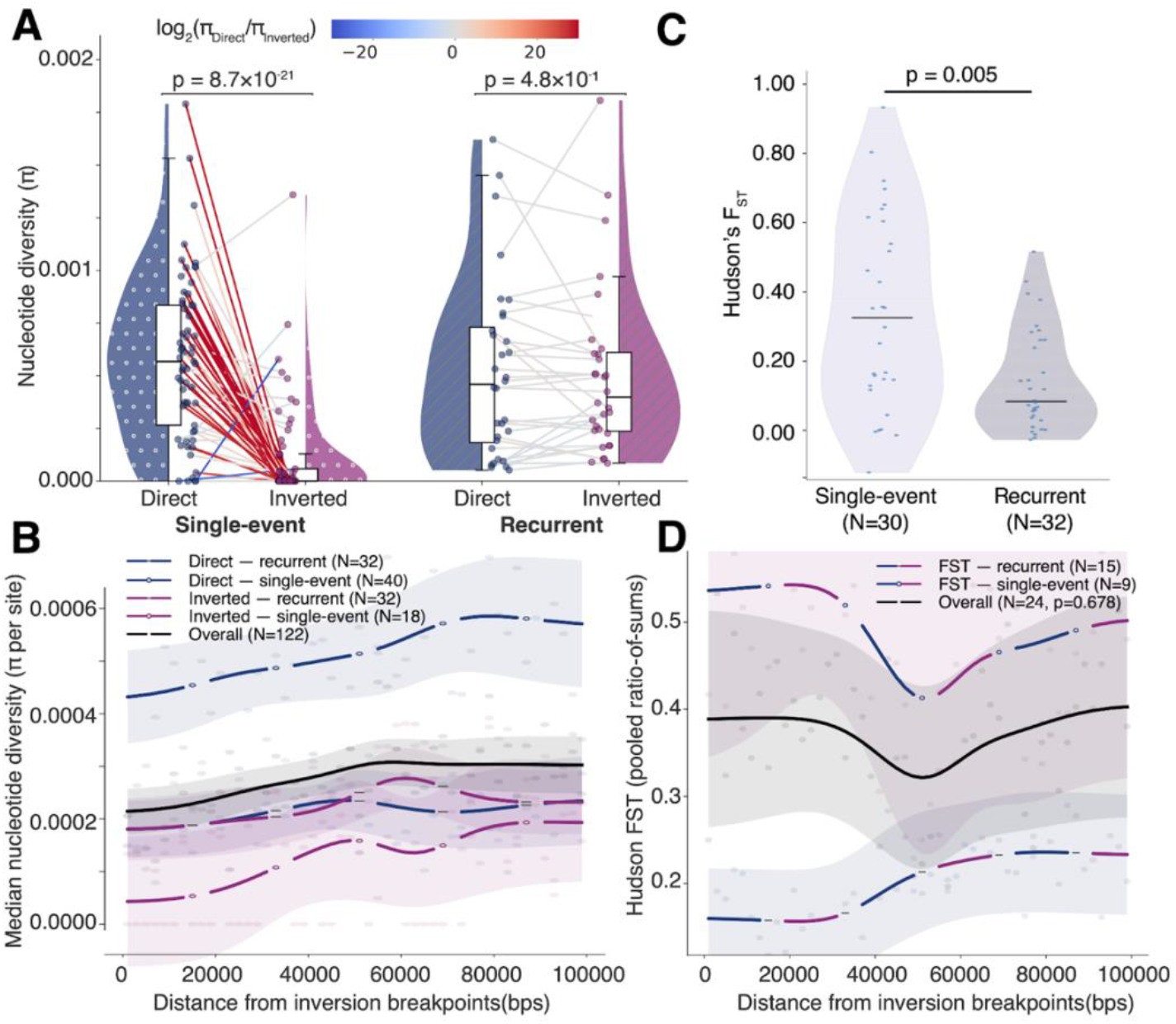
Stratification of genetic diversity (π) and differentiation (F_ST_) across inversion haplotype by recurrence and/or orientation. (**A**) Violin plots show π distributions by inversion event (single-event vs. recurrent) and haplotype orientation (direct vs. inverted). Points are individual π values, where paired lines connect orientations from the same inversion, colored by fold change, log_2_(π_Direct_/π_Inverted_). P values (two-sided Wilcoxon signed-rank) compare orientations within single-event and recurrent groups. Each boxplot represents the median (the thick line in the box), the upper and lower quartiles (depicted by the box), and the 1.5× interquartile range (shown by the whiskers). (**B**) Nucleotide diversity (π) as a function of distance from breakpoints (0) to the first 100,000 base pairs of inversion loci. Points represent the median π calculated from 2 kbp windows using haplotypes within single-event, combined, or recurrent inversion loci. Lines show the smoothed overall trend (Gaussian kernel regression, **Methods**), and the shaded band indicates the ± 1 standard error of the mean calculated per-bin. (**C**) Differentiation between inverted and direct-oriented haplotypes measured by Hudson’s F_ST_ among single-event and recurrent inversion loci. Each point is an inversion locus. P value is computed using a two-sided Mann–Whitney U test. (**D**) Hudson’s F_ST_ as a function of distance from breakpoints (0) to the first 100,000 base pairs within inversion loci. Points represent the pooled *F*_*ST*_ calculated from 2kbp windows using haplotypes within single-event, combined, or recurrent inversion loci. Lines show the smoothed overall trend (Gaussian kernel regression, **Methods**), and the shaded band indicates the ± 1 standard error of the mean calculated per-bin.

Empirical evidence suggested reduced recombination suppression away from the breakpoints of inversions in *Arabidopsis*^28^, with most crossover suppression within 10 thousand base pairs of the inversion’s edge. To test this hypothesis, we assessed the relationship between nucleotide diversity and inversion breakpoint proximity using two different approaches. First, nucleotide diversity was compared between 10 thousand base pairs (10 kbp) breakpoint-flanking regions at each end within the inversion and the 20 kbp middle segment, which requires inversions with at least 40 kbp in total length. Across 315 haplotype sequences that met the criterion, there was significantly lower nucleotide diversity near the breakpoints compared to the middle of the inversion; the middle region had a mean π = 6.74×10^-5^ higher than the flanking regions (paired permutation test p = 0.0027) (**Fig. S5**). When tested by category, both recurrent inverted and recurrent direct haplotypes had significantly lower nucleotide diversity near breakpoints than in the middle (p=0.0025 and p=0.0001, respectively); however, none of the haplotype groups within the single-event loci showed significant differences (**Fig. S6**). As a second approach, we measured π in windows (2 kbp) across the first 100 kbp of each inversion with a total length greater than 100 kbp to obtain a finer resolution across individual loci (N=24). We chose to focus on the first 100 kbp to ensure that enough windows were measured while keeping as many inversions in different sizes as possible (N=122). We found that overall nucleotide diversity near breakpoints is significantly lower compared to sequences away from the breakpoints (ρ=0.525 p<0.001) (**Fig. 2B**). In addition, reduced π near breakpoints was observed across all categories, with the lowest diversity near breakpoints found in single-event inverted haplotypes (N=18, ρ=0.485, p<0.001) (**Fig. 2B**), and the highest in direct haplotypes of recurrent inversions (N=32, ρ=0.315, p=0.026). Together, consistent with observations in other organisms, our results support the widespread phenomenon of reduced recombination suppression away from inversion breakpoints.

Suppressed recombination is expected to create genetic barriers, and in turn result in genetic isolation between orientations in the inverted locus. To test this hypothesis, we studied genetic differentiation between categories of haplotypes, stratified by orientation and recurrence. We first computed the *F*_*ST*_ statistic to assess patterns of differentiation between inverted and direct haplotypes at each inversion locus in our sample, excluding any locus in which there were no polymorphic sites or where there were fewer than two haplotypes for either orientation. Significant differences were found between recurrent and single-event inversions in the amount of genetic differentiation between the direct and inverted haplotypes (**Fig. 2C**). The sequence within a typical recurrent inversion had 3.9-fold lower differentiation (median *F*_*ST*_ = 0.083, n = 32) compared to single-event inversions (median *F*_*ST*_ = 0.33, n = 30) (p = 0.005, Mann-Whitney U test). This suggests that haplotype toggling at the recurrent inversions periodically removes genetic barriers between heterokaryotypes and allows gene flow to occur. In addition, because recombination suppression does not uniformly affect the entire inversion locus, we expect genetic differentiation between orientations to vary across the inversion sequence. Our *F*_*ST*_ analysis showed that measurements near breakpoints are often higher than regions away from them (**Fig. 2D**). However, such a pattern is only found inconsistently across inversions, and appears to be more substantial in single-event inversion loci than in recurrent inversions, consistent with the expectation of stronger genetic barriers around breakpoints between inverted and direct oriented haplotypes (**Fig. 2D**). Overall, our results indicate substantial differentiation between orientations, with 37 inversion loci with *F*_*ST*_ > 0.2 and 12 with *F*_*ST*_ > 0.5, demonstrating complex patterns of evolutionary forces at inversion loci.

### Differential conservation between recurrent and single-event inversions

Inversions can affect the conservation of protein-coding sequences (CDS) through reduced recombination and selection. Weaker selection is expected, due to reduced effective population size, in these haplotypes and can lead to less effective purging of deleterious variants. However, reduced recombination at inversions can also prevent beneficial alleles from being shuffled away and in turn can facilitate selective sweeps. On the other hand, recurrent inversions may retain genetic diversity (**Fig. 2A**) due to haplotype toggling resulting in restoring recombination. The interplay between suppressed recombination and selection at inversion loci is complex, and it is unclear whether selection acts consistently across genes in different inversion categories. Here, we examined two specific questions: are coding sequences within inversion loci conserved across inversion categories, and is there any evidence showing differentiation in genes within inversions?

To test the first question, we analyzed protein-coding sequences (CDSs) within inversion loci. For each CDS, we quantified conservation as the fraction of identical sequences across all pairs of inverted haplotypes and, separately, across direct haplotypes. We grouped these conservation metric values into four haplotype categories (single-event-direct, single-event-inverted, recurrent-direct, recurrent-inverted), and tested for differences in this proportion between all categories (6 tests in total). In the single-event inverted category, 102 CDSs were excluded due to their associated inversion loci having only a single inverted haplotype. For each CDS, we modeled the proportion of identical sequences using a binomial generalized linear model, with terms for the effect of recurrence status, the effect of haplotype orientation, the interaction between recurrence status and orientation, CDS length, the number of haplotypes included for the CDS, and inversion size. To account for non-independence across CDSs within each inversion, we used cluster-robust standard errors grouped by inversion (**Methods**). Our model, weighted so that each inversion contributed equally, estimated (via marginal means) that the single-event inverted group had the highest proportion of identical sequences, at 99.9% identical, followed by the recurrent inverted group (81.8%), then the recurrent direct group (81.4%), and the lowest being the single-event direct group (77.4%). Indeed, the single-event inverted haplotype group had a significantly higher proportion of identical CDSs compared to each of the three other categories (Benjamini–Hochberg [BH] p = 9.2×10^-7^ to 1.5×10^-6^), with no other comparisons reaching significance (**Fig. 3A**). This indicates that inverted haplotypes of single-event inversions almost always contain identical CDS and thus have remarkably conserved protein-coding sequences, compared to the much lower conservation observed in the inverted haplotypes of recurrent inversions.

**Fig. 3.**
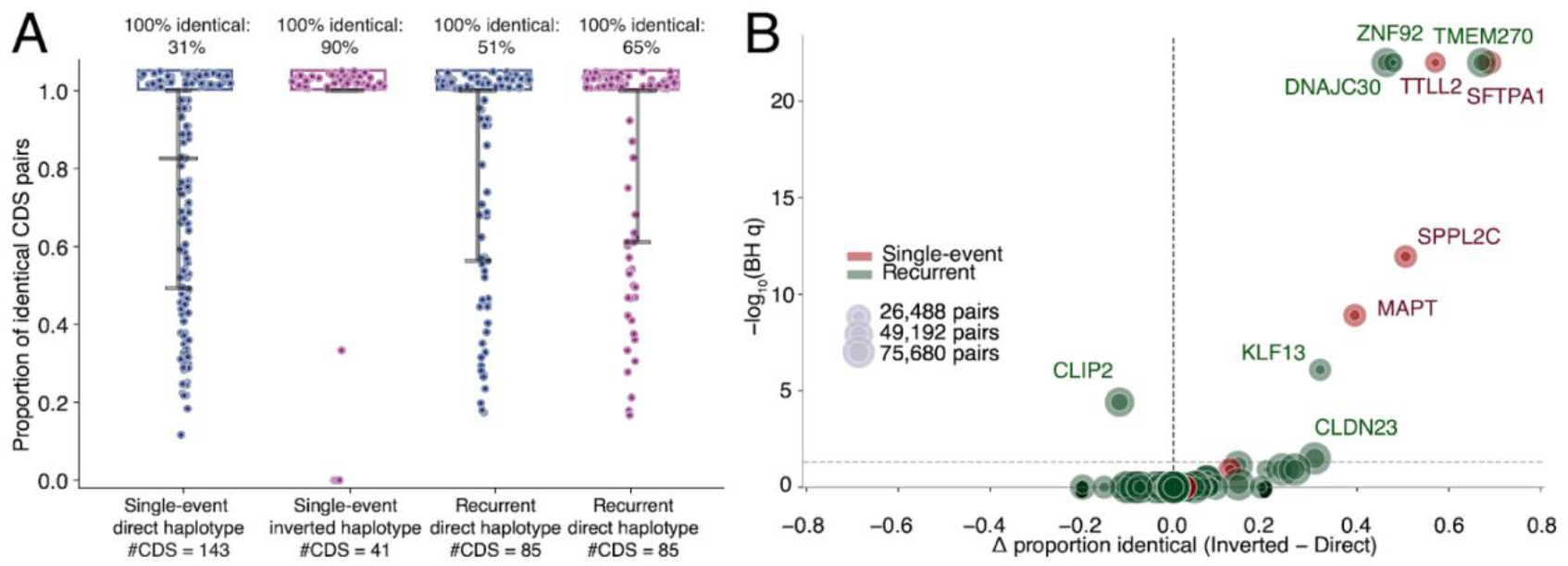
Protein coding sequence (CDS) conservation across inversion loci and categories. CDS conservation is defined as the fraction of identical CDS found among all pairs within each haplotype category. (**A**) Distributions of CDS conservation across the haplotype categories. Each point is a CDS, and the vertical line represents the interquartile range, with upper and lower brackets indicating Q1 and Q3. The horizontal line represents the median. (**B**) A volcano plot shows differential genes (x-axis) and significance (y-axis) in conservation between inverted and direct haplotypes, stratified by inversion recurrence (red: single-event inversion locus, green: recurrent inversion locus). Each circle represents a CDS with its size scaled with the number of sequence pairs. Significance of the differentiation is computed using a generalized linear model with corrected p values using the Benjamin-Hockberg procedure.

To test differentiation in CDS conservation between inverted and direct haplotypes, we designed a test to assess any significant excess or reduction in CDS conservation in the inverted haplotype group (**Methods**). In order to account for the dependency among pairwise observations due to the fact that each haplotype affected multiple CDS pairs, we used leave-one-haplotype-out jackknife standard errors (**Methods**). Using this method, we identified 10 genes showing significant differences (BH q < 0.05) in the fraction of completely identical CDS between inverted and direct haplotypes (**Fig. 3B**); of which, four and six are within single-event and recurrent inversion loci (**Table S2**). Interestingly, the direction of effect was not uniform across these CDSs: in the *CLIP2* gene, inverted haplotypes of the recurrent inversion at chr7:73,113,989-74,799,029 had 11.8% fewer identical coding sequences compared to direct haplotypes (**Fig. 3B**). For all other significant CDSs (*DNAJC30, SFTPA1, TMEM270, TTLL2, ZNF92, SPPL2C, MAPT, KLF13, and CLDN23*), inverted haplotypes had a significantly greater proportion of identical sequences, ranging from 30.8% to 68.9% (BH p values from 0.032 to <1×10^-22^). In the *MAPT* gene, for example, 100% of the pairs from the inverted haplotypes were identical compared to only 60.5% of direct sequence pairs (BH p = 1.25×10^-9^, **Fig. 3B)**.

In addition, we examined fixed nucleotide differences for individual CDS between orientations. After excluding loci with fewer than three inverted haplotypes, we found that eight CDSs carry fixed differences between orientations. Notably, five of these are located in the 17q21 inversion locus, including *MAPT* with eight sites fixed (**Fig. S7**) and *SPPL2C* with seven sites fixed. Together, our results show marked differences in protein-coding sequences at both recurrent and single-event inversion loci, with the inverted haplotypes showing higher conservation but also fixed differences between orientations, suggesting actions of positive selection.

### Selective Pressure on Genes Within Inversions

To assess the possibility of genic evolutionary differences between inversion orientations, we computed *dN*/*dS* along each CDS within inversion loci (**Methods**). We tested for protein-coding evolution differences between inverted and direct haplotypes by comparing the likelihood of two models (Yang 2007) for each gene. The null was constructed via a single *dN*/*dS* estimate shared for all human branches. For the alternative model, sites in the inverted and direct group are free to have different *dN*/*dS* (ω). Trees were constructed over the entire inversion locus sequence using IQ-TREE^29^ with a chimpanzee alignment (*Pan troglodytes*, panTro5) used to root the tree^30^.

After QC of the CDS sequences (**Methods**), we evaluated 34 genes for differences in *dN*/*dS* between inversion orientations. Six CDSs were significant in the clade model after BH-correction. In five out of the six CDSs, the inverted haplotype clade had higher *dN*/*dS* in the codons predicted to differ between orientations. Using the estimates from the alternative model, *MUC20*, found in a 264-kbp inversion on chr3:195,692,124-195,956,606, showed variants with higher *dN*/*dS* in inverted haplotypes (ω=0.22) compared to direct haplotypes (ω=0.00) (BH p = 0.00029). The *CTRB1* gene (inversion locus: chr16:75,206,214-75,222,748) had codons with slightly higher *dN*/*dS* in inverted haplotypes (ω=0.0001) compared to direct haplotypes (ω=0.0) (BH p = 0.00029). The 747 kbp chr7:5,989,046-6,735,643 inversion locus had two genes containing sites with differing *dN*/*dS*: the *USP42* gene (BH p = 0.00583) and the *AIMP2* gene (BH p = 0.0234), where both genes’ inverted haplotype groups have sites with higher *dN*/*dS* compared to the direct haplotypes. For these genes, this indicates that some codons are under a stronger purifying constraint within the direct haplotypes. *GDF10*, in the chr10:46,983,451-47,468,232 inversion locus, similarly showed higher *dN*/*dS* for sites in the inverted sequences (BH p = 0.0173). Uniquely, *ZNF92* in the 313 kbp chr7:65,219,157-65,531,823 inversion was the only gene which had sites with higher *dN*/*dS* for the direct sequences, compared to the inverted sequences (BH p = 0.040). In total, we detect multiple inversion-overlapping coding sequences with evidence of differential selection between inverted and direct orientations, most notably *MUC20, USP42*, and *AIMP2*.

### Disease associations with inversion status

Inversions themselves can have direct effects on carriers through mechanisms, such as disrupting functional sequences by their breakpoints and altering the regulatory landscape^31,32^. To search for possible phenotypic outcomes of inversions, we leveraged data collected in the NIH *All of Us* (AoU) Research Program (v8)^33^ to conduct a series of phenome-wide association studies (PheWASs) (**Methods**). To ensure phenotype quality and relevance, we focused on a total number of 1,096 PheCodes as the test phenotypes after excluding PheCodes that have less than 15% heritability predicted in the UK Biobank, <1,000 AoU cases. We also excluded highly overlapping or correlated phenotypes and restricted to sex-specific samples when ≥99% of cases were of one sex (**Methods**). We imputed inversion allele dosage in AoU participants using a partial least squares regression trained on SNVs within inversion loci identified in the 82 phased haplotypes from Porubsky et al. (2022)^5^ (**Methods**). Imputation performance is assessed using a 5-fold cross-validation procedure and a likelihood ratio test comparing against a constant predictor (**Methods**). We found that 18 out of the 93 inversions have reliable imputation performance (r^2^ > 0.3 and BH p < 0.05, likelihood ratio test) (**Fig.S8**). To increase the test power, we further removed inversions that show less differentiation between inverted and direct-oriented haplotypes based on Hudson’s *F*_*ST*_ (*F*_*ST*_ < 0.5), which led to a final set of nine inversions, all from single-event inversions (**Fig. 2B**). Finally, our PheWAS was conducted using a logistic regression model, controlling for age, age^2^, sex, 16 genetic principal components to account for population structure, and predicted genetic ancestry categories provided by the *All of Us* program^33^.

Using our approaches, we found four of the nine inversions carrying at least one significant signal (false discovery rate = 0.05, BH correction) (**Fig. 4**). A 49-kbp inversion on chromosome 6 (chr6:76,109,081-76,158,474, **Figs. 4A & 5A**) was associated with increased risk for *E. coli* infections at OR = 1.80 (CI: 1.385-1.919, BH p = 0.041) and sepsis (OR = 1.50, CI: 1.28-1.56, BH p = 0.013). The ICD codes for the *E. coli* phecode include various subtypes of *E. coli* infection, including, for example, pneumonia due to *E. coli*, enteropathogenic *E. coli* infection, and *E. coli* as the cause of diseases classified elsewhere. While this inversion is intergenic, SNVs at this locus have been reported and associated with femur total bone mineral density via interactions with the gut-microbiota genus *Lactococcus* ^34^. In addition, we found a 675-kbp inversion on chromosome 10 (chr10:79,542,901-80,217,413, **Figs. 4B** and **5A**) associated with a positive DNA test for human papillomavirus subtypes linked to cervical cancer (OR = 1.233, CI: 1.12-1.35, BH p = 0.0064) and a 19-kbp inversion on chromosome 12 (chr12:46,896,694-46,915,975, **Figs. 4C and 5A**) associated with conjunctivitis (pink eye) (BH p = 0.00035), inflammation of the eye (BH p = 0.0015), acne (BH p = 0.0032), migraine (BH p = 0.048), disorder of nervous system (BH p = 0.025), and epidermal thickening (BH p = 0.043), with ORs ranging from 1.088 to 1.168.

**Fig. 4:**
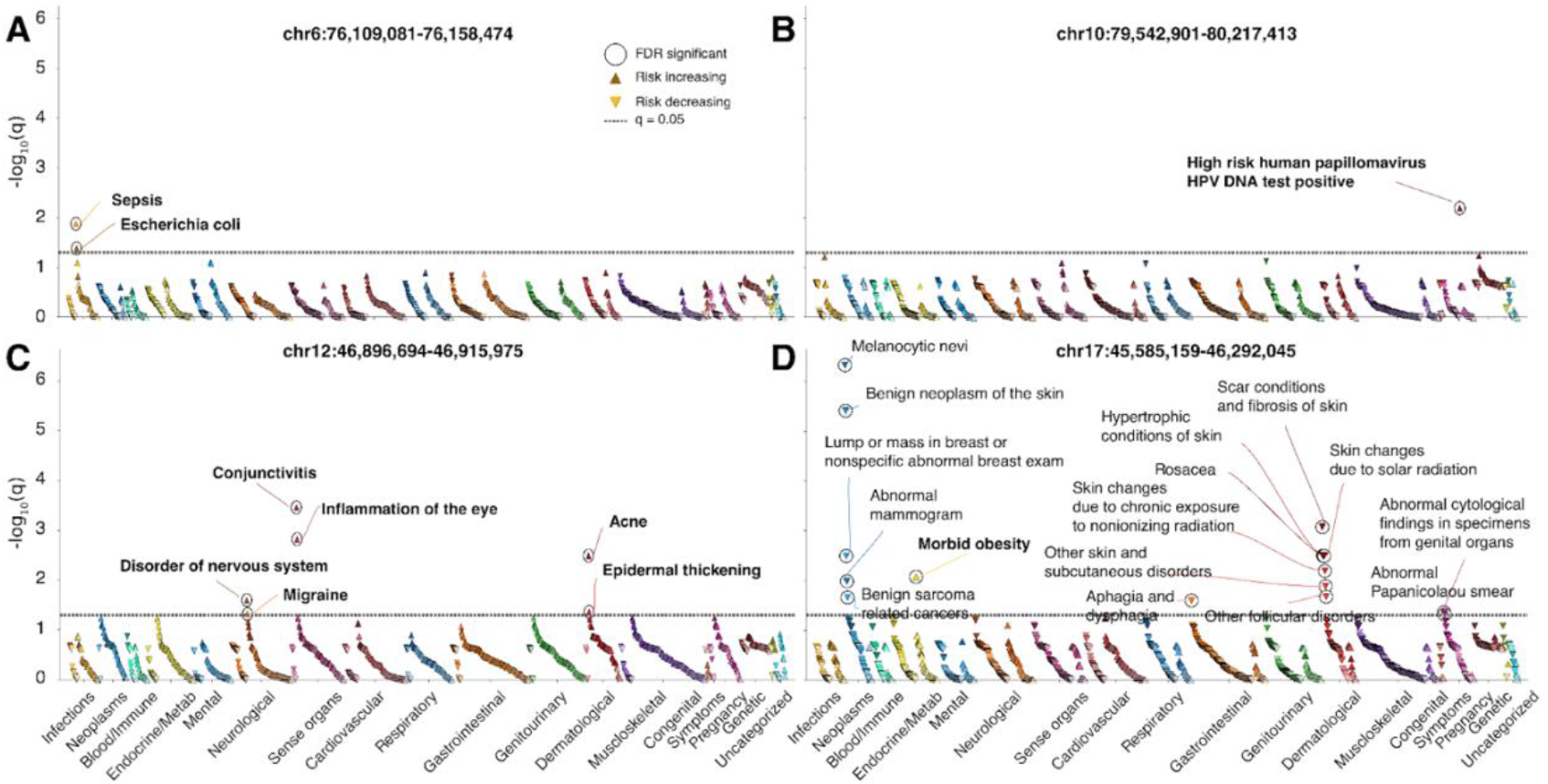
Phenome-wide inversion association in the NIH *All of Us* research cohort (v8). Each Manhattan plot shows significant associations between phenotypes (phecodes) and the imputed inversion genotype dosage (**Methods**). (**A**) A 49 kbp inversion at chr6:76,109,081-76,158,474. (**B**) A 67 kbp inversion at chr10:79,542,901-80,217,413. (**C**) A 19 kbp inversion at chr12:46,896,694-46,915,975. (**D**) A 706 kbp inversion at chr17:45,585,159-46,292,045. Each triangle is a phecode. Upward and downward triangles represent increasing and decreasing risk, respectively. Each circle indicates a significant inversion-phecode association (FDR (q) < 0.05). Dotted lines are the phenome-wide FDR significance level at 5%, across all tested phecodes and nine inverted loci. Each bolded phecode indicates an association with an increased risk.

The strongest signal of association is located at chromosome 17q21 (chr17:45,585,159-46,292,045). Our analysis showed that while the 706-kbp inversion allele is associated with higher risk for morbid obesity (OR = 1.076, CI: 1.04-1.11, BH p = 0.0084), it is associated with decreased risk of multiple PheCode conditions (**Figs. 4D & 5A**). Notably, this is the same inversion where we observed significantly different CDS conservation and fixed nucleotide differences between orientation in multiple genes, including *SPPL2C* and *MAPT* (**Fig. S7**). Among the PheCodes showing reduced risks, a variety of skin conditions are significantly associated with the inverted allele (OR range=0.875-0.942, **Fig. 5A**), including melanocytic nevi (moles), benign neoplasm of the skin, skin changes due to solar radiation, rosacea, hypertrophic conditions of skin, and scar conditions and fibrosis of the skin. In addition, the inverted allele is associated with lower risk of multiple gynecologic traits, including “lump or mass in the breast or a nonspecific abnormal breast” exam (OR = 0.928, CI: 0.9-0.96, BH p = 0.0032), abnormal mammogram (OR = 0.905, CI: 0.897-0.939, BH p = 0.010), and abnormal Papanicolaou smear (OR = 0.898, CI: 0.88-0.94, BH p = 0.048). Other PheCodes associated with the 17q21 inverted allele show reduced risk, including benign sarcoma-related cancers (OR = 0.936, CI: 0.93-0.961, BH p = 0.022) as well as aphagia and dysphagia (inability or difficulty with swallowing) (OR = 0.928, CI: 0.918-0.957, BH p = 0.025).

**Fig. 5.**
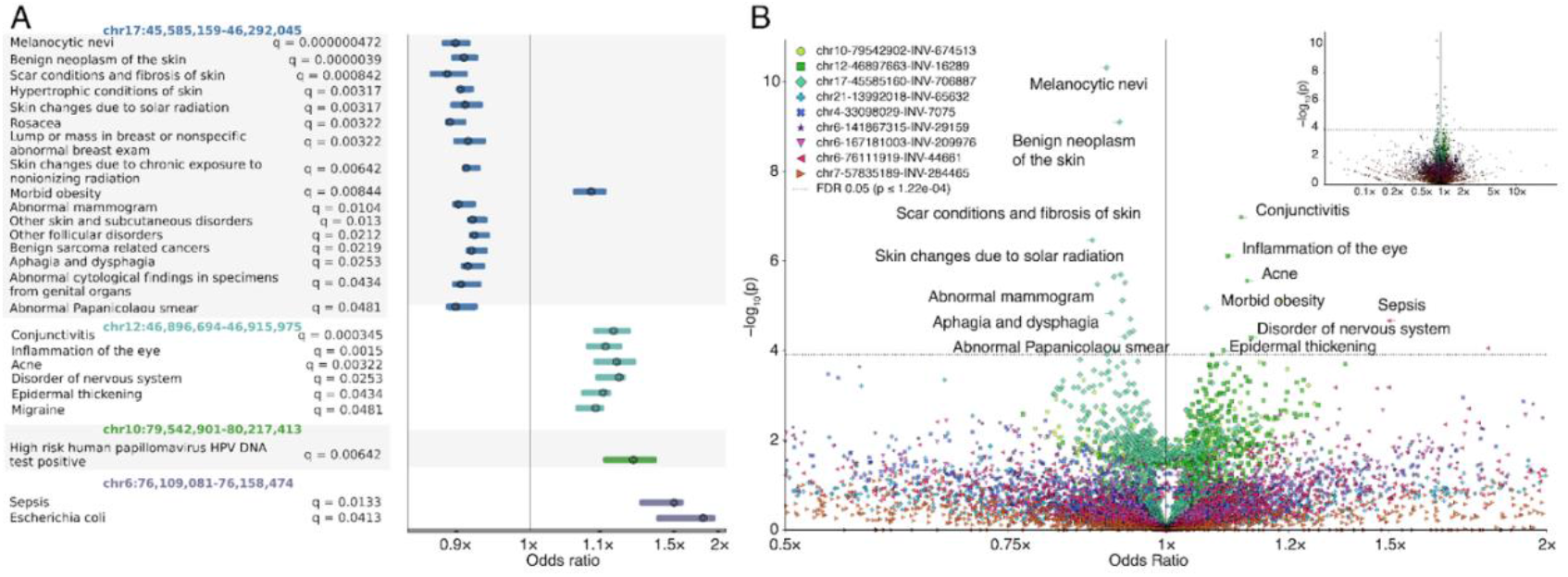
Effect estimates of the inversion on phecodes. (**A**) A forest plot illustrates the significant associations between individual inversions (n=4) and phenotypes (phecodes) in the PheWAS analysis. The x-axis represents the odds ratios (OR) with their corresponding 95% confidence intervals, while the y-axis lists the significant PheCodes. Q-values are calculated using the Benjamini-Hochberg false discovery rate (FDR) method. (**B**) A volcano plot highlights differential effects (measured by odds ratio) across all 1,096 PheCodes and nine high-quality, imputable inversions in the PheWAS analysis. The x-axis shows the log-transformed odds ratios, and the y-axis represents the Benjamini-Hochberg-adjusted p-values on a –log_10_ scale. The horizontal dashed line indicates the phenome-wide significance threshold across all inversions. Each point represents an inversion–phecode association test. The main plot displays an x-axis range of OR = 0.5 to OR = 2.0, while the inset plot shows the full x-axis scale.

Multiple association studies have reported signals from this 17q21 locus associated with breast cancer, cognitive decline^35^, heart failure^36^, and obesity^37^. To further assess concordance of our PheWAS disease associations at the 17q21 inversion locus, we analyzed family history survey data, regardless of personal history or ICD code status. In addition to breast cancer and obesity, we included cognitive decline (dementia, memory loss, or impairment) and heart failure as we observed nominal associations in mild cognitive impairment (BH p = 0.057, OR = 0.855), diastolic heart failure (BH p = 0.057, OR = 1.115), and malignant neoplasm of the breast (BH p = 0.064, OR = 0.901). To attain comparable odds ratios, we modeled the expected allele count for each relative category (children, siblings, parents, and grandparents) as expected under Mendelian inheritance given the participant’s own inversion dosage (**Methods**). Our analysis of family history showed that three of the four diseases had overlapping confidence intervals and the same direction of effect with the main PheWAS results, including breast cancer: OR = 0.91 (CI: 0.859-0.964, p=0.001), cognitive decline: OR = 0.919 (CI: 0.866-0.974, p=0.005), and obesity: OR = 1.097 (CI: 1.04-1.157, p=0.001), but was insignificant for heart failure: OR = 1.043 (CI: 0.984-1.106, p=0.154). Our PheWAS and family history analyses suggest that the 17q21 inversion allele protects against breast cancer and cognitive decline but increases obesity risk. Alongside reported positive selection signals^38^, the present evidence for significant differentiation in CDS conservation and fixed nucleotide differences in coding sequences between inversion orientations support the theory that inversions can enhance fitness by capturing multiple advantageous alleles while reducing the efficiency of purging weakly deleterious variants due to suppressed recombination^16^.

## Discussion

Despite the important role of inversions in biology, studying the evolution and function of inversions has been proven difficult, not only due to the difficulty of resolving their sequences, but also due to complex, often interacting processes affecting their evolution. Our understanding of human inversions is particularly limited because they are frequently flanked by long and highly identical segmental duplications, making the detection of such mutations challenging, leaving a gap in knowledge of human variation. In this study, we overcame this challenge by leveraging the published high-quality, sequenced-resolved inversion callset from Porubsky et al. (2022)^5^. This allows us to systematically interrogate different aspects of human inversions to study their evolution and roles in human biology for the first time. Using large scale structural coalescent simulations, we were able to demonstrate the statistical power for distinguishing recurrent inversions from single-event ones following the evolutionary framework established by Porubsky et al. (2022)^5^, providing the basis for further dissecting the evolution of human inversions in a refined scale.

Our work demonstrates that in humans, recurrent inversions operate under a fundamentally different evolutionary regime. We find that single-event inversions create stronger genetic barriers, sharply reduce diversity within inverted haplotypes, whereas recurrent inversions only maintain modest barriers and preserve gene flow via periodic recombination. As a result, recurrent direct and inverted haplotypes have similar diversity, while single-event inverted haplotypes show much lower diversity. Notably, classic population genetics theory predicts that gene flux resulting from recombination between inversion arrangements is most suppressed near breakpoints, leading to greater divergence there than in the inversion center – a suspension bridge pattern^3^. While showing strong divergence near breakpoints, our results also show additional peaks moving toward the center in both single-event and recurrent inversions. Such a pattern might arise due to locally adapted alleles associated with the inversion driving differentiation between orientations^39^ or reduced flux at linked neutral sites due to selection against deleterious variants exchanged between heterokaryotypes resulting in reduced gene flux at linked neutral sites^16^. Because the same patterns can arise from distinct mechanisms that are not easily distinguishable, this highlights the complex interplay among different evolutionary forces at inversions.

Interestingly, the 17q21 inversion locus (chr17:45,585,159-46,292,045) is mostly associated with decreased risk for associated diseases, whereas the inversion locus chr12:46,896,694-46,915,975 is mostly associated with increased risk for diseases. It is not clear why an inversion allele, originally only present in a single individual, would increase in frequency despite being associated with increased risk for disease: there are multiple potential explanations. For example, it is possible that the inversion is nearly neutral or deleterious, yet brought to higher frequency due to drift, or it is possible that the inversion has beneficial effects unmeasured by data available in the biobank or by phecode definitions. It should be noted that the inverted allele (H2) in the 17q21 inversion is the ancestral allele, which arose over 2 million years ago and was subsequently nearly replaced by the derived, direct (H1) allele^7,14,40^.

The 17q21 inversion allele is unique for having clear pleiotropic effect in this analysis, being associated with higher risk for morbid obesity, lower risk for antecedents to breast cancer and family history of breast cancer, and lower risk for a variety of skin conditions, most of which are not known to be commonly comorbid with each other. Moles/benign skin neoplasms, rosacea, hypertrophic skin conditions/scar conditions, and solar-related skin changes are four distinct conditions of the skin. Darker skin is associated with higher hypertrophic scar and keloid formation^41,42^, whereas lighter skin is related to skin changes due to solar radiation and melanocytic nevi^43,44^. Since the 17q21 inversion allele is associated with decreased risk for each of these conditions, it is unlikely that confounding due to residual population substructure is the sole explanation for this cluster of associations.

Given that *MAPT*, a gene encoding a protein critical to Alzheimer’s disease pathogenesis, is within the 17q21 inversion locus, it is particularly interesting that there are two suggestive signals associating the 17q21 inversion allele with Alzheimer’s disease-related phenotypes. The main PheWAS found a risk-decreasing association (BH p = 0.057) for mild cognitive impairment, and similarly, family history was suggestive of reduced risk of mild cognitive impairment (p = 0.055). It is plausible that decreased risk for a variety of prevalent diseases affects human fitness; therefore, these results provide a potential explanation for the positive selection on the 17q21 inverted (H2) allele observed in the modern day by Stefansson et al. (2005)^38^. The wide array of associations with the 17q21 inversion could be due to supergene mechanics, direct transcriptional or other regulatory impacts of inverted synteny, or linked variants affecting genes within the region.

Due to the widespread disease associations of inversions, understanding inversion population genetics and disease associations could improve disease risk prediction and lead to mechanistic understandings relevant to clinical disease interpretation. Future work may assess disease association at a wider variety of inversions, particularly those with evidence of differential conservation or selection between orientations, or with algorithmically predicted transcriptional impact. Pangenomic analysis methods may increase the feasibility of incorporating inversions, which often lie in structurally complex regions, into polygenic risk scores. As reference panels increase in accuracy and number of samples, and as biobanks gain more long-read data, we expect more inversion–disease associations, due to both improved inversion imputation power and the ability to directly call the inversions. Our imputation method is able to be used on microarray data in addition to short-read whole-genome sequencing data, allowing for the possibility of increasing sample size with additional PheWAS analysis in the *All of Us* and UK biobank microarray cohorts. Many inversions, especially single-event inversions, are not common, appearing in only a few haplotypes in the dataset used. Future pangenomic research, with larger cohort sizes, may be able to identify a larger number of genomic inversions. This analysis only considers balanced inversions, in which there are no associated insertions or deletions, and further restricts analysis to only inversions in which recurrence can be confidently called.

Overall, we produce the first genome-wide analysis of the sequence content within a high-quality callset of human inversions and discover multiple inversion–disease associations with impact to human health. We characterize differences in diversity, differentiation, conservation, and selection, showing that recurrent inversions are generally different from single-event inversions, with lower differentiation between orientations and higher diversity. We conclude that inversions have critical relevance to common diseases and have impactful evolutionary consequences, especially on diversity and differentiation.

## Methods

### Simulations for power analysis of recurrent inversion detection

To model recurrent inversions, we developed a process leveraging structured coalescent with limited to no gene flow (**Fig. 1A and 1B**). When an inversion event occurs, recombination between heterokaryotypes is largely suppressed due to lack of homologous pairing at the locus during meiosis. This results in a localized barrier limiting the exchange of genetic material between the two haplotypes. Such a process can be modeled under the structured coalescent framework, where an inversion event creates a subpopulation (e.g., inverted haplotypes) diverged from the ancestral population (e.g., directly oriented haplotype) and isolated from each other by eliminating any gene flow between them (**Fig. 1A**). Recurrent inversions are therefore modeled as any subsequent divergence events in the same process (**Fig. 1B**). Each divergence introduces a bottleneck (90% reduction) in the new subpopulation of different orientation from its parental population). Gene flow is restored between any subpopulations that are in the same haplotype orientation as recombination is allowed and genetic variation can be shuffled among them. Finally, inverted and direct haplotypes are sampled at pre-defined frequencies f_inv_ and 1-f_inv_, respectively, from mixtures of populations with the same orientation at a random rate.

We modeled four different time depths of inversion events and simulated data using msprime^45^. For single-event models, inversions were set to occur at 500, 250, 100, and 50 kya. For three-recurrent inversion models, we considered four scenarios with events at (i) 500, 250, 100 kya, (ii) 250, 100, 50 kya, (iii) 100, 50, 25 kya, and (iv) 50, 25, 10 kya. Simulations were run under recombination rates of 1×10^-10^ (low), 1×10^-8^ (moderate), and 1×10^-6^ (high) per base pair per generation. Each locus was 200 kbp in length, with an ancestral effective population size of 10,000 and a generation time of 25 years. We used a mutation rate of 1.25×10^-8^ per base pair per generation and allowed gene flow of 1×10^-8^ per chromosome per generation between haplotypes of the same orientation. Code for generating simulations and recurrent inversion inference is available at https://github.com/hsiehphLab/inversionSimulation.

### Variant data filtering

Polymorphic sites were retained if they were biallelic single nucleotide variants and if they had a genotype quality (GQ) above 30 for all samples. In addition, variants within centromeres, acrocentric arms, known gaps, and segmental duplications with sequence identity > 99% (GRCh38) are removed to ensure the quality of variant genotypes.

### Nucleotide diversity

Nucleotide diversity (π) was calculated for each defined group of haplotypes, over each genomic inversion locus, with an effective sequence length adjusted for masked subregions. All unique pairs of haplotypes within each orientation group were used to determine the number of nucleotide differences. π for the group was computed via the sum of differences divided by the number of sites compared. π could not be computed when there were fewer than two haplotypes in the group.

The model for computing the recurrence effect on orientation has a per-locus outcome of the log difference between the inverted and direct nucleotide diversity (with an epsilon floor at half of the 1st percentile of positive nucleotide diversity values to avoid taking the log of zero). We regress the per-locus log difference between orientations on recurrence status. The model specification is as follows:

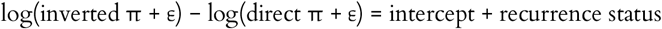

Since the outcome is per-locus, the model removes the overall locus baseline, making the recurrence term coefficient equal to the difference in the orientation effect between recurrent and single-event groups, in order to detect orientation-by-recurrence interaction.

### Distance from breakpoints

Per-site nucleotide diversity (π) was calculated from phased haplotype genotype data. For each inversion locus, per-site π was calculated separately within the inverted haplotype group and within the direct haplotype group. Loci under 100 kbp were excluded, and positions greater than 100 kbp away from the nearest breakpoint were not considered. The relationship between mean π per bin and the distance from the segment edge was evaluated using Spearman’s rank correlation.

### Coding sequences

Coding sequence (CDS) information was derived from a gene annotation file in GTF format from GENCODE (Release 47, GRCh38.p14, “[b]asic gene annotation”). For each distinct gene (gene_id) identified on the chromosome, a single representative transcript was selected to define the CDS structure used in subsequent analyses. Transcripts were first ranked based on annotation tags present in the GTF attributes, following a predefined priority order: MANE_Select, MANE_Plus_Clinical, CCDS, appris_principal_1, GENCODE_Primary, Ensembl_canonical, and basic (listed from highest to lowest priority). If multiple transcripts for the same gene shared the highest identified priority level, the tie was resolved by selecting the transcript with the greatest cumulative CDS length, calculated as the sum of the lengths of all its individual CDS segments.

These sequences were retained only if the CDS segments overlapped an inversion locus. Sequences were constructed after variant quality filtering, such that low-quality variants were replaced by the reference genome’s allele at that position, in order to filter low-quality data while retaining unbroken sequences. CDSs were excluded if the total length of the sequence was not divisible by three, if the sequence did not begin with the ATG start codon, or if there was an internal stop codon (TAA, TAG, TGA) in the coding frame.

For the comparison of sequence conservation by category, coding sequences with fewer than two sequences were excluded. For each CDS, the outcome is the number of identical sequence pairs out of the total number of pairwise comparisons in that CDS. This is modeled as a binomial, with the denominator equal to the total number of pairwise comparisons, and a mean equal to the probability that a randomly chosen pair within that alignment is identical. Each coding sequence contributes proportionally to the number of pairwise comparisons to prevent over-weighting of very rare inversion alleles. To account for multiple CDSs overlapping the same inversion locus, we used a sandwich covariance estimator (Huber– White) with clustering by inversion identity for standard error (SE), which inflates SE to be robust to arbitrary covariance structure for CDSs within an inversion. The model testing proportional sequences is specified as follows.

#### Unadjusted for covariates

logit(probability that a random pair of CDS sequences is identical) = intercept + indicator of recurrent status + indicator of inverted haplotype group + indicator of recurrent status × indicator of inverted haplotype group

#### Including additional covariates

logit(probability that a random pair of CDS sequences is identical) = intercept + indicator of recurrent status + indicator of inverted haplotype group + indicator of recurrent status × indicator of inverted haplotype group + ln(CDS length in base pairs) + ln(inversion length in base pairs) + ln(number of haplotypes in the CDS)

### dN/dS Calculation and Analysis

To estimate selective pressure on protein-coding genes, we estimated the ratio of nonsynonymous to synonymous substitution rates per coding sequence, including both direct and inverted sequences. The calculation was performed using the “codeml” program from the PAML software package. Coding sequences were excluded if the median human-chimp divergence was greater than 10% (due to misalignment risk), if there were fewer than 3 haplotypes in each group, or fewer than 2 variable codons. Coding sequences were further excluded if the tree had fewer than 4 taxa, or if there was not at least one pure internal node (whose descendants are all direct or all inverted) for each orientation (to ensure the tree topology is informative). For each coding sequence, we executed “codeml” using the clade model C setting (runmode = 0) and the F3×4 codon frequency model (CodonFreq = 2). For the null model, any codon’s dN/dS is free to be indicative of neutral, positive, or purifying selection, but dN/dS is constrained to a single estimate at each codon across all haplotypes, both direct and inverted. Benjamini–Hochberg correction was applied across all likelihood ratio tests. The dN/dS estimates are from the full model in which dN/dS is allowed to vary between groups. These dN/dS estimates are specifically for the subset of codons which are predicted to differ between orientations, as opposed to a single estimate across the entire coding sequence.

### Inversion Imputation

We trained imputation models for each inversion locus. These models predict the expected genotype dosage of the inverted allele given local SNP dosages. For each inversion, we used Partial Least Squares (PLS) to associate SNP dosage patterns within 50 kbp of the inversion breakpoints as well as within the inversion, to the observed inversion dosage in the phased dataset from Porubsky et al. (2022)^5^. We included synthetic diploid genomes in the training data by combining different haplotypes that were not seen in the original data. For example, any two samples have four haplotypes total, but only two observed diploid genomes. Since there are ten unique possible combinations of these haplotypes, an additional eight diploid genomes can be constructed. This approach does not require phased data during inference.

Per-inversion imputation significance was tested by comparison to a simple model which only had an intercept term, and always predicted a constant value equal to the mean inversion dosage seen during training, in order to verify that the model was not merely learning inversion frequency. The number of components in each inversion model was chosen by cross-validation to minimize mean squared error. The final model is refit on all samples in addition to the synthetic data. Predicted inversion dosage = intercept + (component 1 contribution) + (component 2 contribution) + … + (component K contribution), where each component is equal to (slope for component k) × (weighted sum of the allele counts). After training, this model reduces to an intercept plus one coefficient per SNP, times that SNP’s allele count. Weights were chosen via the partial least-squares regression objective maximizing the covariance between the weighted sum of SNP counts and the current residual outcome (that is, the residual outcome after accounting for all previously fit components). We required at least 20 samples with inversion dosage information to proceed with imputation training. The number of PLS components to include was determined via up to 3-fold cross-validation. An up to 5-fold cross-validation was used to estimate out-of-sample performance.

### PheCode filtering and QC

To test for phenotypic associations, we used data from the NIH *All of Us* Research Program cohort version v8. The analysis was conducted using a logistic regression model, controlling for age, age^2^, sex, 16 genetic principal components to account for population structure, and genetic ancestry categories. Statistical significance was determined using a false discovery rate of 5% to correct for multiple testing.

SNPs were only considered for inclusion if they were biallelic, had a 100% call rate among phased samples from Porubsky et al. 2022^5^, and were also found in the *All of Us* ACAF (Allele Count/Allele Frequency threshold callset) short-read whole genome sequencing (srWGS) dataset. We define SNPs within the ACAF set as those with a call rate above 95% in the *All of Us* v8 cohort ACAF PLINK 1.9 files. For inversion inclusion, we required an imputation accuracy of r^2^ > 0.3, required significantly lower absolute error compared to a constant, intercept-only predictor (Wilcoxon one-sided p < 0.05), and required most of the average genetic difference between orientation to be explainable by between-orientation diversity (that is, *F*_*ST*_ > 0.5).

We used a phecodeX^46^ map to provide mappings between PheCodes and ICD codes, and Pan-UKBB^47^ to provide mappings between ICD codes and UKBB phenotype identifiers (phenocodes) as well as phenocode heritability. PheCodes were associated with phenocodes if there was at least one ICD code shared between them. We consider PheCodes for further analysis if there is no SNP heritability data available, or, if there is evidence of non-zero liability-scale SNP heritability for at least one associated phenocode (ACAT across ancestry groups, one-sided z-test, Benjamini–Hochberg across UKBB phenocodes) in the Pan-UKBB data, and when estimated SNP heritability combined across groups ≥ 0.15 (estimated by a precision-weighted linear random-effects model). For each phenocode and population, standard error of heritability, between-population variance, between-phenocode variance, and residual variance were used to weight each heritability estimation observation via restricted maximum-likelihood estimation. This approach estimates the precision-weighted mean heritability for a phecode, across all associated phenocodes and populations.

### Phenotypic Association Analysis

Participants were defined as cases for the phecode if they had at least one ICD-9 or ICD-10 code mapped. We also require at least 1,000 cases for the phecode in the *All of Us* cohort. Controls were individuals without any ICD code from the entire phecode category associated with the phecode, to avoid contamination of related or comorbid phenotypes. PheCodes with over 90,000 cases were excluded due to being overly general (e.g., viral infection). A phenotype was excluded if 70% or higher of its cases were shared with another phenotype, or if it had a binary Pearson coefficient greater than 0.7 with another phenotype. If 99% or more cases were one sex, we restricted the samples under consideration to that sex only and omitted the sex covariate. 1,096 final PheCodes were tested.

We tested the association between SV dosage and each eligible phecode using logistic regression with a logit link and a likelihood-ratio test. In the model, we include the imputed expected SV dosage, age at the end date of observation, inferred sex (derived from *All of Us* DRAGEN ploidy calls), and the first 16 principal components (precomputed by *All of Us*) for ancestry main effects (adjusting for prevalence differences varying along the genetic ancestry continuum). We report the overall odds ratio (OR) per unit of dosage. All individuals were unrelated. Zero-variance covariates (e.g., inferred sex) are removed per-phecode. For each phecode, we compute an LRT for the contribution of the imputed SV dosage term.

This compares the full model to a model without SV dosage. Across tested phenotypes, we apply Benjamini–Hochberg (BH) (α=0.05) correction. The PheWAS model is specified as follows: logit(case status) = intercept + imputed inversion dosage + genetically inferred sex + 16 PCs + age + age^2^ + year of birth + genetic ancestry categories (“afr,” “amr,” “mid,” “sas,” “eur”)

For each BH-significant locus, we assessed ancestry-specific effect heterogeneity by testing a dosage × ancestry term via LRT. Ancestry groups were pre-computed by *All of Us* and inferred via random forest on genetic principal components. Here, we compare a full model to the model omitting this interaction term. Reported confidence intervals are derived from profile-likelihood, due to its increased robustness to non-normality compared to standard Wald confidence intervals. If profile-likelihood did not converge, Wald confidence intervals were reported instead.

Family history for the 17q21 association analyses was assessed from *All of Us* “Personal and Family Health History” survey questions. For breast cancer, obesity, heart failure, dementia, and memory loss or impairment, participants answered “who in your family has had” the condition. Possible options include self, mother, father, sibling, daughter, son, or grandparent. “Self” is not used in this follow-up analysis since many of the same individuals were present in the main PheWAS already. We computed the cohort allele frequency as the mean of the participants’ inversion dosages. For each family tier, we used the relatedness coefficient r_g_ multiplied by the dosage of the related participant to derive an expected allele count, assuming additive inheritance.

Intercepts and age interactions with family relationship allow the baseline level and its dependence on age and cohort to differ by family relationship, while the genetic effect is constrained to a single shared coefficient across tiers. The model outcome is if the participant has at least one affected relative in a given tier. Outcome = intercept + r_g_ * (participant’s inversion dosage) + four family tier indicator terms + age + age2 + year of birth + four family tier indicators * age + family tier indicators * age2 + family tier indicators * year of birth + sex + 16 PCs + indicator terms for categorical genetic ancestry groups (“afr,” “amr,” “mid,” “sas,” “eur”). Age was defined by participant age at the end of observation.

## Funding

P.H. is supported by an NIH Pathway to Independence Award (NHGRI, 5R00HG011041).

## Author contributions

S.C.S. and P.H. designed and planned experiments. S.C.S. B.B., and P.H. analyzed sequencing data and performed bioinformatics analyses. S.C.S. and P.H. designed and performed population and statistical genetics inferences and analyses. S.C.S. and P.H. wrote the manuscript.

## Competing interests

S.C.S is a co-founder of PolyCypher Health, Inc. The rest of the authors declare no competing interests.

